# A Transient Intermediate Populated in Prion Folding Leads to Domain Swapping

**DOI:** 10.1101/724666

**Authors:** Balaka Mondal, Govardhan Reddy

## Abstract

Aggregation of misfolded prion proteins causes fatal neurodegenerative disorders in both humans and animals. There is an extensive effort to identify the elusive aggregation-prone conformations (*N*\*) of prions, which are early stage precursors to aggregation. Ve studied temperature and force induced unfolding of the structured C-terminal domain of mouse (moPrP) and human prion proteins (hPrP) using molecular dynamics simulations and coarse-grained protein models. Ve find that these proteins sparsely populate intermediate states bearing the features of *N*\* and readily undergo domain-swapped dimerization by swapping the short *β*-strands present at the beginning of the C-terminal domain. Structure of the *N*\* state is similar for both moPrP and hPrP, indicating a common pathogenic precursor across diferent species. Interestingly, disease-resistant hPrP (G127V) showed a drastic reduction in the population of *N*\* state further hinting a pathogenic connection to these partially denatured conformations. This study proposes a plausible runaway domain swapping mechanism to describe the onset of prion aggregation.

## Introduction

Aggregation of prion proteins in the central nervous system leads to transmissible spongiform encephalopathies (TSE), a diverse group of lethal neurodegenerative disorders also known as prion diseases(*1*). According to the protein only hypothesis(*1, 2*), a misfolded form of prion protein known as the pathogenic scrapie form (PrP^SC^) autocatalyzes the conversion of a normal cellular prion protein (PrP^C^) to the toxic scrapie form. Experiments(*3*) infer that PrP^SC^ is an oligomer containing approximately 12-24 misfolded prion monomers chemically identical to PrP^C^(*4*), except with a higher *β*-sheet content(*5*). The mechanism for structural transition from PrP^C^ to PrP^SC^ is ambiguous and the structure of the protein in the PrP^SC^ form is elusive. An important question in prion aggregation is what are the early structural changes in PrP^C^ that facilitates the formation of PrP^SC^ oligomers and fibrils.

The structure of membrane-bound cellular mouse prion protein (moPrP) is not available. Recombinant moPrP (without glycosylation and GPI anchor), whose three dimensional structure is resolved, serves as a reference structure for cellular moPrP (Figure 1A)(*6*). The N-terminal region of the moPrP lacks structure while the C-terminal domain has a globular fold, which is majorly *α*-helical with three *α*-helices (*α*_1_: Asp144 to Met1,54, *α*_2_: Gln172 to Thr193 and *α*_3_: Glu200 to Tyr226) and a short antiparallel *β* sheet (*β*_1_: Met129 to Gly131, *β*_2_: Val161 to Tyr163) (Figure 1A)(*6*). The prion monomer undergoes structural transition as it aggregates to form distinct oligomeric assemblies and amyloid fibrils. The long incubation time for prion disease indicates that the conversion of a PrP^C^ monomer to the PrP^SC^ form by itself must be thermodynamically unfavorable at physiological conditions and must be at least initially aided by the interaction of PrP^C^ with other monomers or cofactors to lower the thermodynamic barriers for oligomer formation(*7*).

**Figure 1:**
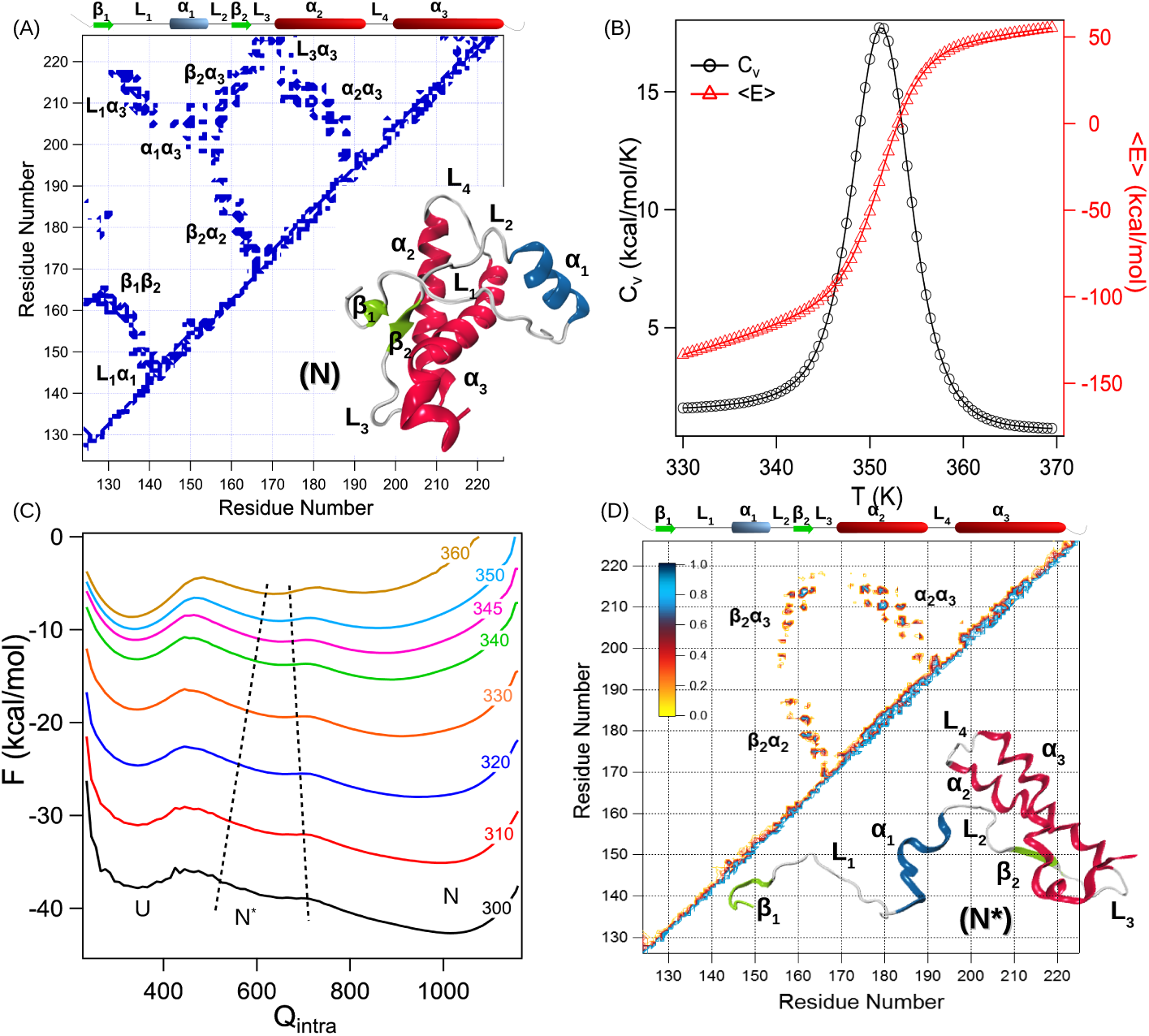
Structure and stability of recombinant moPrP. (A) Cartoon representation and contact map of structured C-terminal domain of moPrP from Gly124 to Tyr226 (Protein Data Bank (PDB) ID: 1AG2). In the native state, the protein has three *α*-helices (*α*_1_ − *α*_3_), two *β*-strands (*β*_1_ − *β*_2_) and loops (*L*_1_ − *L*_4_) connecting diferent SSEs. The contact map shows important tertiary contacts between diferent SSEs. (B) Average potential energy (⟨*E*⟩) and specific heat (*C*_*v*_) plotted as a function of temperature (*T*), show two state folding behavior without any intermediates. (C) Folding free energy (*F*) projected onto the number of native contacts present in the protein monomer (*Q*_*intra*_) is shown at diferent temperatures. A shallow minima corresponding to an aggregation-prone intermediate state (*N*\*) is observed in addition to the folded (*N*) and unfolded (*U*) states. The *N*\* state appears at *Q*_*intra*_ ≈ 500 − 700. (D) Representative structure and contact map of the *N*\* state. Contacts are missing in the *N*\* state due to the unfolding and detachment of *β*_1_ − *L*_1_ − *α*_1_ segment. Structural perturbations are also observed in the folded *β*_2_ − *α*_2_ − *α*_3_ sub-domain, where the loop *L*_3_ connecting *β*_2_ and *α*_2_, and C-terminal end residues of *α*_3_ helix (residues Lys220 to Tyr226) undergo partial unfolding leading to the rupture of contacts between *α*_3_ and *L*_3_.

There have been extensive efforts using a range of experimental and computational techniques to understand the relative stabilities of various secondary structural elements (SSEs) present in prions((*8–11*)), and the effects of temperature(*11, 12*), force(*13, 14*), salt(*15*), pH((*16–24*)), mutations(*8, 25–36*), disulfide bond reducing agents((*37–40*)), salt bridges(*8, 41*) and protein-protein interactions((*28, 42–44*)) on the conformational transitions in prions during the initial stages of aggregation. The protein is dynamic and rapidly undergoes conformational transitions between an ensemble of structures(*45*). A number of aggregation prone partially folded intermediates (*N*\*) were also identified both experimentally(*46–51*) and computationally(*9, 13, 44, 52*). Some common features of these intermediate states include a disordered *α*_1_ and *β*_1_ region, with *α*_2_ helix converted to *β*-sheet(*53*) (Figure 1A). Experiments((*54–57*)) further showed that the highly hydrophilic *α*_1_ helix is very much flexible and can get detached from the protein core stimulating aggregation while not getting converted to a *β* sheet(*8*). Using computations, Dima and Thirumalai(*8, 58*) have predicted that *α*_2_ helix is frustrated due to the presence of a patch of residues T188VTTTT193, which are highly conserved in mammalian prion proteins and have a high *β*-sheet formation propensity. Mutations, which stabilize the *α*_2_ helix are shown to either prevent or slow down the oligomer formation(*59*) confirming the predictions, which are also subsequently verified in a number of other experiments((*59–63*)). Identifying aggregation prone transient *N*\* states, which are excitations above the native folded monomers((*64–68*)), are extremely important to prevent aggregation. Despite significant progress, a clear demonstration of a prion intermediate driving the initial stages of oligomerization is still elusive.

Using molecular dynamics simulations and coarse-grained self-organized polymer-side chain (SOP-SC) protein model(*69, 70*) of mouse(*6*) and human prion proteins(*71*) we show that these proteins sparsely populate two intermediates in their folding pathway. One of the intermediates is an aggregation-prone (*N*\*) state, which readily dimerizes revealing the early stages of prion aggregation, and the second intermediate is a short lived metastable (I) state(*51, 72*). In the *N*\* state, the SSEs *β*_1_, *L*_1_ and *α*_1_ are detached from the folded protein core comprised of *β*_2_, *α*_2_ and *α*_3_. In the initial stages of dimerization, the disordered segments (*β*_1_-*L*_1_-*α*_1_) in *N*\* interact with the other protein in solution leading to the formation of a domain-swapped dimer. This allows us to speculate that the major conformational transitions in the structured core should occur in the late stages of dimerization as *α*_2_ and *α*_3_ helices are converted to *β*-sheets as predicted by the previous computations(*8, 58*) and the parallel in-register *β* sheet model(*60, 73*). Interestingly, we further find that the frequency of populating both the *N*\* and *I* states in the disease resistant G127V mutant of human prion protein (hPrP) is an order of magnitude lower compared to the wild type. The mechanism of *N*\* dimerization revealed in this work can be the harbinger of prion oligomerization leading to the formation of PrP^SC^ oligomers and finally mature fibrils. The predictions from the simulations can be further verified by experiments.

## Materials and Methods

Molecular dynamics simulations are carried out using coarse-grained SOP-SC model(*69, 70*) where each amino acid residue is represented by two beads. One bead represents the backbone atoms and the second bead represents the side chain atoms of an amino acid. The SOP-SC model for moPrP is constructed using the NMR structure(*6*) with PDB ID: 1AG2, and hPrP is constructed using the NMR structures(*71*) with PDB ID: ,5YJ4 and ,5YJ,5. The energy of a protein conformation in the SOP-SC model given by a set of coordinates {**r**} is,

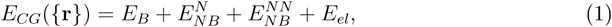

where *E*_*B*_ is the energy due to bonds between the beads, 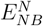 and 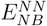 are the energy terms due to non-bonded native and non-native interactions, respectively, and *E*_*el*_ is the energy due to electrostatic interactions present between charged residues in the protein. To compute the folding thermodynamic properties we performed low friction Langevin dynamics simulations. Equilibrium properties like average internal energy ⟨*E*⟩, specific heat capacity 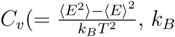, is the Boltzmann constant) and folding free energy *F* projected onto the number of native contacts *Q* is computed using the simulation data and weighted histogram analysis method (WHAM)(*74*). Brownian dynamics(*75*) simulations are performed at *T* = 300 K to study prion dimerization and force induced unfolding. A symmetric Go-type potential(*76, 77*) was used to construct the dimer energy function (Figure S9). Detailed description of the simulation methodology, SOP-SC model and symmetric Go model with associated energy functions and parameters are given in the SI.

The total number of native contacts in a protein conformation is given by 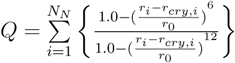, where *N*_*N*_ is the total number of native contacts, *r*_*cry,i*_ and *r*_*i*_ is the distance between the *i*^*th*^ pair of beads in the native structure and in a conformation given by {**r**}, respectively, and *r*_0_ is the cut-of distance. Radius of gyration is computed using 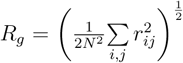, where *N* is the total number of coarse-grained beads, and *r*_*ij*_ is the distance between beads *i* and *j*. End-to-end distance *R*_*ee*_ is the distance between the backbone beads of the N and C termini residues.

## Results and Discussion

### *N*\* State is Populated in the moPrP Unfoiding Pathway

To compute the folding thermodynamics of the structured C-terminal domain of moPrP (residues Gly124 to Tyr226), we performed low friction Langevin dynamics simulations using coarse-grained SOP-SC model(*69, 70*). The SOP-SC model is constructed using the solution NMR structure (PDB ID: 1AG2)(*6*). Detailed description of the coarse-grained model and simulation methodology is in the supplementary information (SI). The disulfide bond between Cys179 and Cys214, which holds *α*_2_ and *α*_3_ helices together is disabled as experimental studies(*37–40*) have shown that aggregation is accelerated when the disulfide bond is reduced. The computed average internal energy (⟨*E*⟩), and heat capacity (*C*_*v*_) as a function of temperature (*T*) show that folding from the unfolded (U) to the native (N) state occurs cooperatively in a two-state manner in agreement with the experiments(*50, 78, 79*) (Figure 1B). The melting temperature (*T*_*M*_) of the protein obtained from the simulations is ≈ 3,50 K, which is in reasonable agreement with the experimentally measured value of 338 K(*79*). In contrast to *C*_*v*_, the folding free energy (*F*) projected onto the number of native contacts present in the protein (*Q*_*intra*_) show evidence for the population of an aggregation prone intermediate state *N*\* in the folding pathway (Figure 1C).

At *T*_*M*_, the free energy diference between *N* and *N*\* state is 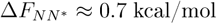 with an energy barrier 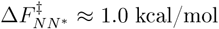. The major barrier separating *N* from the *U* state is 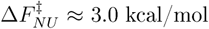. Lower value of 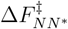 compared to 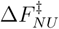 suggests that *N*\* state can be directly populated from the native state ensemble without crossing over the major energy barrier. The contact map of the *N*\* state shows that the SSEs *β*_1_, *α*_1_ and *L*_1_ are detached from the folded protein core and are disordered, while the rest of the protein core composed of *β*_2_ − *α*_2_ − *α*_3_ remains folded (Figure 1D). Contacts present between *β*_1_ and *β*_2_, *L*_1_ and *α*_1_, *L*_1_ and *α*_3_, and, *α*_1_ and *α*_3_ are missing in the *N*\* state confirming the complete unfolding of the *β*_1_ − *L*_1_ − *α*_1_ segment. The folded core also exhibits structural perturbations in the *L*_3_ loop region connecting *β*_2_ and *α*_2_, and the C-terminal end of *α*_3_ helix as evident from the contact map (Figure 1D,S1). A very similar intermediate state with disordered *β*_1_, *L*_1_ and C-terminal end of *α*_3_ has been reported in moPrP experiments performed at pH 4(*51*). Population of prion folding intermediates with some of the structural characteristics observed in *N*\* are also reported in several other experiments(*24, 35, 46, 47, 49, 50, 55, 57*). Structural features of the *N*\* state with the exposed *β*_1_ strand is compatible to form steric zipper conformations and can initiate aggregation(*80*).

We performed Brownian dynamics simulations to probe the force-induced unfolding pathways of moPrP (see SI for details). The backbone bead of the N-terminal residue Gly124 is fixed and a constant force *f* is applied on the backbone bead of the C-terminal residue Tyr226. For *f* = 4,5 pN, we spawned 100 unfolding trajectories at *T* = 300 K. The plot of end-to-end distance *R*_*ee*_ as a function of simulation time shows that the *N*\* state is observed in the unfolding pathways (Figure 2A) and is populated in all the 100 independent trajectories. The *R*_*ee*_ corresponding to the *N*\* state is ≈ 140 Å. The time dependent variation in the values of radius of gyration *R*_*g*_ and *Q*_*intra*_ also show the presence of *N*\* in the unfolding pathway (Figure S2). To monitor the unfolding mechanism, we studied the progression of rupture of tertiary interactions between the various SSEs by computing the fraction of native contacts between the SSEs (*f*_*ss*_) as a function of time (Figure 2B). In the folded state, tertiary contacts between the following SSEs are present in the prion protein: *β*_1_*β*_2_, *L*_1_*α*_3_, *β*_2_*α*_2_, *β*_2_*α*_3_ and *α*_2_*α*_3_ (Figure 1A). The time-dependent variation in *f*_*ss*_ between diferent SSEs in the unfolding pathway shows that initially *β*_1_ detaches from the *β*_2_ strand followed by the rupture of contacts between *α*_3_ and *L*_1_. This constitutes the *N*\* state, and the contact map shows that its structure with an exposed *β*_1_ strand is very similar to the *N*\* state observed in the temperature dependent unfolding of moPrP (Figure S3). The *N*\* state further unfolds as the remaining contacts between the SSEs *β*_2_, *α*_2_ and *α*_3_ break down simultaneously (Figure 2B,C). This observation is in agreement with the experimental(*50, 81*) findings, which suggest that the transition state consists of native like interactions between *β*_2_, *α*_2_ and *α*_3_ during prion unfolding.

**Figure 2:**
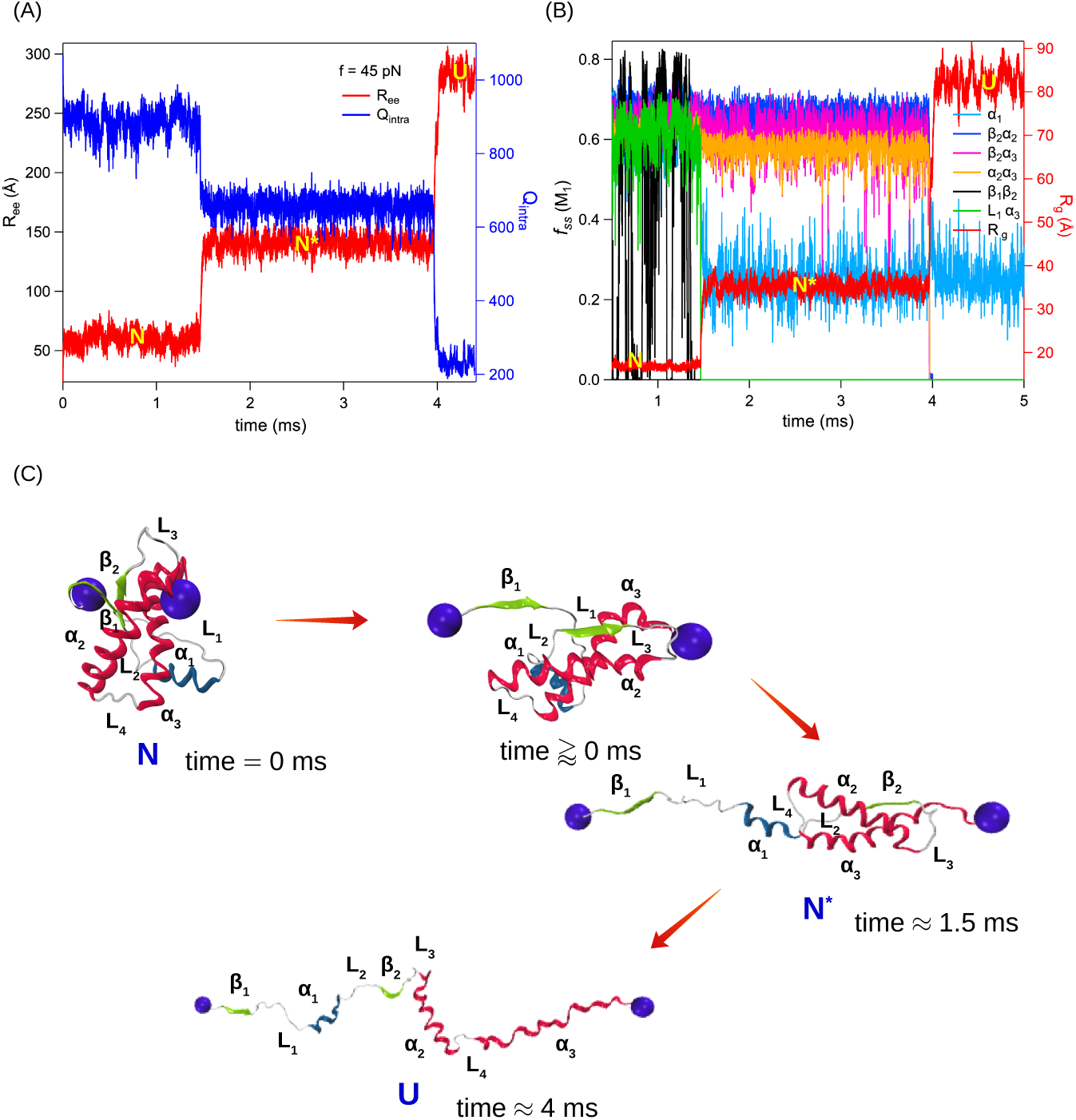
Force induced unfolding of moPrP at *T* = 300 K and *f* = 45 pN. (A) *R*_*ee*_ and *Q*_*intra*_ is plotted as a function of simulation time for a representative unfolding trajectory. The trajectory reveal a three state unfolding pathway with the presence of an intermediate state (*N*\*) at *R*_*ee*_ ≈ 140 Å and *Q*_*intra*_ ≈ 6,50. (B) Fraction of native contacts (*f*_*ss*_) present in diferent SSEs and *R*_*g*_ are plotted as a function of simulation time to monitor the unfolding pathway. *f*_*ss*_ and *R*_*g*_ are plotted on the left and right-axis, respectively. The contacts present between *β*_1_ − *β*_2_ strands break followed by the rupture of *L*_1_ − *α*_3_ contacts populating the *N*\* state. At this stage, *β*_1_ − *L*_1_ − *α*_1_ segment gets detached from the folded sub-domain comprised of *β*_2_ − *α*_2_ − *α*_3_. (C) Representative protein structures from the native state (N), a conformation where the *β*_1_-strand starts to unfold, the *N*\* state and the fully unfolded state (U) are shown.

The unperturbed tertiary contacts between *α*_2_ and *α*_3_ helices in *N*\* led us to the question whether the reduction of disulfide bond between Cys179 and Cys214 is essential for the initial stages of prion misfolding. To verify this we performed both Langevin dynamics and force-induced unfolding simulations of moPrP with an intact disulfide bond (modeled using FENE potential, details in SI). *N*\* state is also populated (Figure S4) in these simulations indicating that reduction of disulfide bond might not be essential for the initial stages of prion misfolding but it can be important in the later stages when major conformational changes take place in the structured core *β*_2_ − *α*_2_ − *α*_3_.

### Propensity of Populating *N*\* State is Low in Prions with Disease Resistant Mutations

Mutant G127V (with M129 polymorphic variant) of hPrP is shown to be disease-resistant(*82*). The mechanism for disease-resistance is unknown, which makes this protein an excellent model system to probe the role of intermediates (if any) in aggregation. To study the folding thermodynamics, we carried out Langevin dynamics simulations of both the disease-resistant (PDB ID: 5YJ4) and wild-type (PDB ID: 5YJ5) hPrPs using coarse-grained SOP-SC model constructed from the solution NMR structures(*71*) determined at pH 4.,5. G127V mutation enhanced the stability of the folded state as the *T*_*M*_ of the protective mutant increased by ≈ 10 K compared to the wild-type protein (Figure S5A,B). A previous study(*71*) indicated that in the mutated protein, *α*_1_ helix gets closer to *α*_2_ and *α*_3_ helices as compared to the wild type, which is evident from the contact maps (Figure S5C,D). This results in a more compact structure and higher thermodynamic stability of the mutated hPrP due to enhanced number of contacts between *α*_1_ and *α*_3_, *L*_1_ and *α*_3_, and, *L*_1_ and *α*_1_. Thermodynamic properties such as ⟨*E*⟩ and *C*_*v*_ as a function of *T* show that both the wild-type and G127V mutated hPrPs exhibit two-state folding (Figure S5A,B). However, free energy *F* projected onto *Q*_*intra*_ shows that the wild-type hPrP populates two intermediate states (*I* and *N*\*) in equilibrium with the folded state *N* (Figure 3A,S6). At *T*_*M*_, the free energy diference between *N* and *N*\* in the wild-type protein is 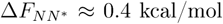 while the energy barrier 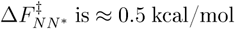 as opposed to a relatively higher barrier 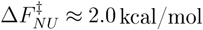 separating *N* and *U*. The state *I* appears as a shoulder in the free energy surface at *Q*_*intra*_ ≈ ,500. A wider native basin and lower energy barriers to the intermediate states in the free energy surface suggest a conformationally flexible native state. Contrastingly, *I* is absent in the free energy surface of the disease-resistant hPrP, while *N*\* shows a much reduced stability and appears as a shoulder (Figure 3B,S7A).

**Figure 3:**
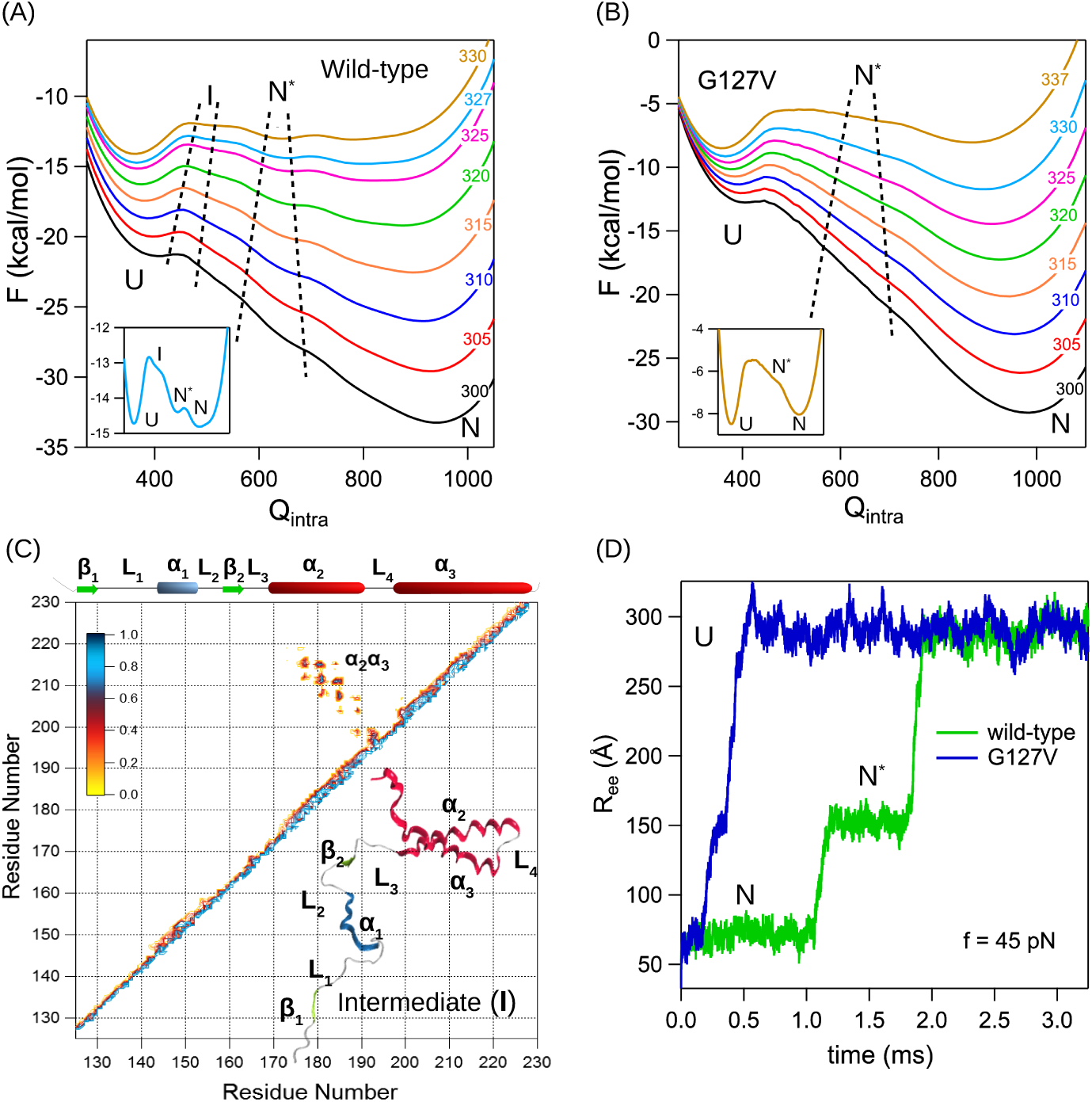
Folding free energy (*F*) of hPrP is projected onto *Q*_*intra*_ for (A) wild-type (PDB ID: ,5YJ,5) and (B) G127V disease-resistant mutant (PDB ID: ,5YJ4). Both the proteins show minima corresponding to the folded state (*N*) and denatured state (*U*) at *Q*_*intra*_ ≈ 900 and 350, respectively. In the case of wild-type hPrP, two intermediate states, *I* and *N*\*, are observed at *Q*_*intra*_ ≈ 500 and ≈ 700, respectively. However, for disease-resistant hPrP, *I* is absent and *N*\* appears as a shoulder due to reduced stability. In the inset of panels-(A) and (B), *F* plot at *T*_*M*_ is shown for clarity. (C) Contact map of the short-lived transient intermediate state (*I*) populated in wild-type hPrP folding. The state *I* consists of a structured sub-domain comprised of *α*_2_ − *α*_3_ helices, and a disordered segment comprised of *β*_1_ − *α*_1_ − *β*_2_. (D) Force induced unfolding of wild-type (in green) and disease-resistant (in blue) hPrPs. *R*_*ee*_ is plotted as a function of time for *f* = 45 pN. A three state unfolding pathway with the population of *N*\* state is confirmed in all the unfolding trajectories of both wild-type and disease-resistant hPrP.

To probe whether the intermediate states also get populated during force induced unfolding of hPrPs, we performed Brownian dynamics simulations of wild-type and disease-resistant hPrPs at a constant force *f*. We spawned 100 independent unfolding trajectories for each protein at *T* = 300 K, where the backbone bead of the N-terminal residue is fixed and a constant force *f* = 4,5 pN is applied on the backbone bead of the C-terminal. Both the proteins populated *N*\* state in their unfolding pathways whose structural features are similar to the intermediate state populated in the moPrP simulations (Figure 3D,S8). However, we did not observe a separate *I* state during the force induced unfolding of both the proteins. Thus *I* might be a high energy short-lived intermediate, which could not be detected in force induced unfolding of hPrPs. Contact map of *N*\* state is similar to that of moPrP (Figure S8), and features a folded core consisting of *β*_2_ − *α*_2_ − *α*_3_, and an unfolded segment comprised of *β*_1_ − *α*_1_. The state *I*, which is absent in disease-resistant hPrP consists of a folded core *α*_2_ − *α*_3_ and an unfolded segment comprised of *β*_1_ − *α*_1_ − *β*_2_ and was observed experimentally in a full-length moPrP at pH 4(*51, 72*), and in a computational study(*9*). The absence of *I* state in moPrP simulations can be due to the different solvent conditions in the experiments and simulations or reduced-level description of the protein. The average lifetime of *N*\* in the disease-resistant hPrP is less by a factor of ≈ 7 compared to the wild-type. The distributions show that the lifetime of *N*\* state in the mutated protein does not exceed 0.4 ms, while in the wild-type the lifetimes exceed 2 ms (Figure S7B). The longer lifetime of the *N*\* state should facilitate further misfolding and aggregation in wild-type compared to the disease-resistant hPrP.

Experiments(*83, 84*) show that proteins with engineered disulfide bonds, which facilitate the population of intermediates with sub-domain separation similar to *I* and *N*\* states undergo aggregation, and disulfide bonds, which hinder the sub-domain separation suppress aggregation of the protein. Several other experimental studies(*33, 34, 51, 85*) also indicated that *α*_1_ helix initiates misfolding by breaking away from *α*_2_*α*_3_ folded segment prior to the conformational changes in the C-terminal domain. Our simulations support this observation as enhanced contacts between *α*_1_ and *α*_2_*α*_3_ helices in the mutated prion compared to the wild-type, impede the detachment of *α*_1_ from the *α*_2_*α*_3_ folded segment leading to the decreased stability of the *N*\* state in the mutated protein compared to the wild-type (Figure 3). Due to the low stability of the *N*\* state in the mutated prion, the time scales for aggregation increase and disease progression can be inhibited. Structural resemblance between the *N*\* states obtained from moPrP and hPrP simulations indicates the possibility of a general pathogenic intermediate across diferent species (Figure S8).

### Dimerization of the *N*\* State

To probe the early events in prion aggregation where monomeric PrP^C^s get converted to PrP^SC^s and form higher order oligomers, we studied the mechanism of dimer formation in moPrP, which is the simplest oligomer possible. The initial simulation setup was designed to mimic the self-replicated conformational conversion of PrP^C^ to PrP^SC^, which consisted of a monomer in the *N*\* state (*M*_1_) and another monomer in the native folded state (*M*_2_). We chose *N*\* over *I* as the former was a common intermediate obtained in both moPrP and hPrP and had a higher stability. In order to facilitate dimerization, the centre of mass distance between the two proteins was restrained at 18 Å (≈ *R*_*g*_ of a monomer) using a harmonic potential with a spring constant, *k* = 2.0 kcal/mol/Å^2^(*86*). We employed symmetric Go-potential(*76, 77*) (Figure S9) to construct the inter contacts (*Q*_*inter*_) present between the monomers (model details are given in the SI). Brownian dynamics simulations are performed at *T* = 300 K to probe the dimerization kinetics. Out of 100 independent simulations, dimerization was observed in 30 trajectories.

We monitored the time evolution of intra contacts (*Q*_*intra*_) present within monomers *M*_1_ (*N*\* state) and *M*_2_ (*N* state), and inter contacts (*Q*_*inter*_) formed between the monomers to track the dimerization process. The simulation trajectories revealed a two step dimerization process, corresponding to the formation of an intermediate state *D*_1_ (*Q*_*inter*_ ∼ 150) and finally the dimer *D*_2_ (*Q*_*inter*_ ∼ 350) (Figure 4A). Structures of *D*_1_ and *D*_2_ bear the hallmarks of domain swapping, which is the most anticipated mechanism of aggregation in globular proteins. In the dimerized state, the two monomers swapped the flexible *β*_1_ strands while *α*_1_ helix acted as a hinge loop.

**Figure 4:**
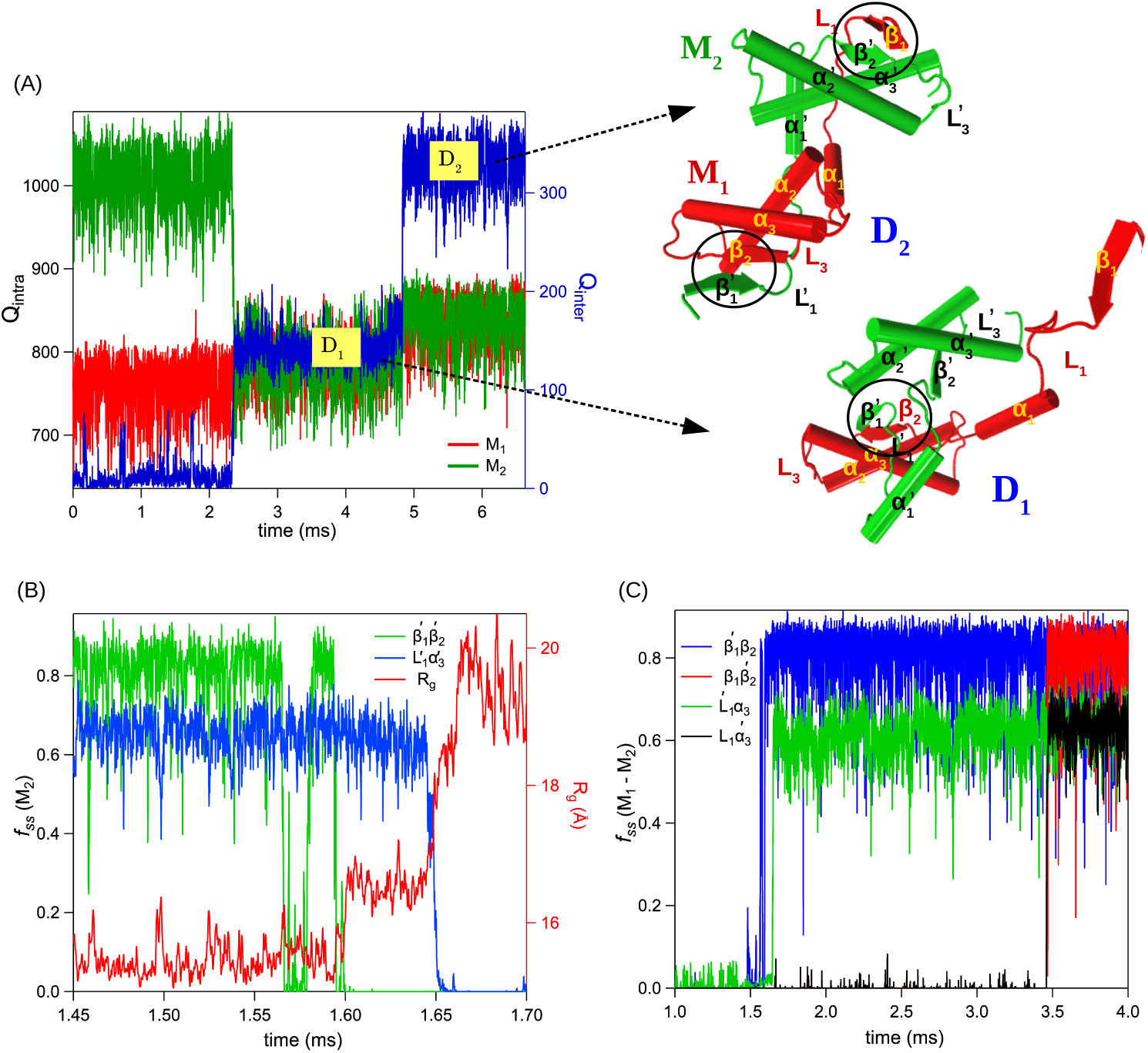
Domain swapped dimerization of moPrP. (A) *Q*_*intra*_ and *Q*_*inter*_ are plotted as a function of time to track the dimerization process. Two states *D*_1_ and *D*_2_ are observed corresponding to *Q*_*inter*_ ≈ 150 and ≈ 350, respectively. Representative structures of *D*_1_ and *D*_2_ states are shown. (B) Time-dependent changes in *R*_*g*_ and fraction of native contacts present in the SSEs of *M*_2_ monomer (*f*_*ss*_(*M*_2_)) during dimerization. Contacts between 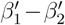 breaks first, followed by the rupture of 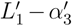 contacts. (C) Time-dependent changes in the fraction of contacts (*f*_*ss*_(*M*_1_, *M*_2_)) between *M*_1_ and *M*_2_ during dimerization. A fully domain swapped dimer (*D*_2_) is obtained when neighboring monomers exchange *β*_1_ − *L*_1_ segment with each other.

The time-dependent changes in *f*_*ss*_ in various SSEs show that the exposed 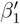 strand in *M*_1_ due to the unfolding of *β*_1_*L*_1_*α*_1_ segment initiates dimerization by forming contacts with *β*1^*!*^ strand in the natively folded monomer *M*_2_, followed by *α*_3_ helix forming contacts with loop 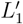 (structure *D*_1_) (Figure 4B,4C,S10A). This disrupts the interactions between the residues in 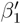 and 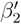, and, 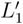 and 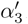. As a result 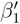 detaches from 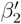, and 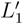 detaches from 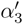, leading to the unfolding of the segment 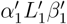 leaving 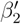 exposed in the *β*_2_*α*_2_*α*_3_ folded core. Finally, the detached *β*_1_ in *M*_1_ forms contact with the exposed 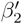 in *M*_2_ to give rise to a fully domain-swapped structure *D*_2_ (Figure 4B,4C,S10B). This suggests a possible runaway domain swapping model(*87*) for prion aggregation, where monomers swap their *β*_1_ strands in end-to-end manner with neighboring partners giving rise to higher order oligomers. In support of our *β*_1_ strand mediated domain swapping model, simulations show that aggregation is initiated by *β*_1_ strand (Figure 4B,C) and its role in pathogenic conversion of prion proteins is also inferred in experiments(*80, 88*). Experimental studies((*88-91*)) show that *β*_1_ strands from human, rabbit and sheep prions form intermolecular *β*-sheet rich oligomers indicating a possible role of *β*_1_ strand in initiating aggregation via steric zipper conformations(*80*).

The domain-swapped dimer structure we obtained from simulations is different from the domain swapped crystal structure of hPrP(*92*). In the dimeric crystal structure, monomers exchange their C-terminal *α*_3_ helices and the disulfide bonds (Cys179-Cys214) originally holding *α*_2_ and *α*_3_ helices in the monomers get reduced. However, a new pair of intermolecular disulfide bonds (Cys179-Cys′214 and Cys′179-Cys214) appear between the swapped *α*_3_ helix from one monomer and *α*_2_ helix from another monomer. Formation of such structure requires reduction of the intramolecular disulfide bonds followed by the formation of inter-molecular disulfide bonds under oxidizing conditions. This domain-swapped structure was observed in a previous computational study(*42*) where dimerization between the two prion monomers was studied with a preformed irreversible intermolecular disulfide bond and a symmetric Go-model potential. The importance of such intermolecular disulfide bonds in prion aggregation is not established yet as it is uncertain whether these bonds are present in prion aggregates(*93, 94*). Moreover, such unique domain swapped structure was observed only in case of wild-type hPrP and not in disease causing pathogenic mutants(*89*). Experiments further show that reduction of disulfide bonds accelerates prion aggregation((*37-40*)), while the hPrP dimer structure crystallized under non-reducing conditions disappears on addition of reducing agents(*92*).

To check the stability of domain-swapped dimers (structure *D*_2_), we converted the coarse-grained domain-swapped dimer structure to a fully atomistic description and ran three independent simulations in the NPT ensemble at *T* = 300 K and *P* = 1 atm (details in the SI). The explicit solvent simulations are performed using NAMD molecular dynamics package(*95*) and CHARMM22 forcefield(*96, 97*).The simulations show that the dimer structure is stable on a timescale of ≈ 700 ns in all the trajectories (Figure S11). In a trajectory, the number of residues forming *β*-strands also increased on the timescale of the simulation indicating that the dimer structure is not an artifact of the coarse-grained model.

Based on the intermediate states and domain swapped dimeric structure obtained in our simulations, we propose a runaway domain swapping model to explain initial stages of aggregation in prion proteins. Aggregation prone *N*\* state (with a detached *β*_1_ *− α*_1_ segment) induces misfolding in native monomeric proteins and form higher order oligomers via swapping of *β*_1_ strands in end-to-end manner. However, our model accounts only for the initial stages of aggregation, which is probably further accompanied by *α*-helix to *β*-sheet conformational changes in the folded core consisting of *β*_2_ *− α*_2_ *− α*_3_ as predicted by Dima and Thirumalai(*8, 58*). Computations (*58*)show that *α*_2_ helix is shown to have a high propensity for *β*-sheet formation and experiments(*59*) support the prediction by showing that oligomerization can be prevented by stabilizing the *α*_2_ helix. Thus although domain swapping of N-terminal *β* strands initiates the aggregation process, the final aggregate originates from the structural conversion in the folded core and can be prevented by stabilizing the secondary structures in the folded core.

## Conclusions

The structural transition of cellular prion proteins to the pathogenic PrP^SC^ form is a complex process involving major conformational transitions. There is growing evidence((*64-68*)) that high energy partially unfolded intermediates play an important role in the early stages of aggregation of globular proteins. In this paper, using coarse-grained SOP-SC model for prions and molecular dynamics simulations, we showed that aggregation prone short-lived metastable intermediate states are populated from the prion folded states in both temperature and force induced unfolding. In these states, the N-terminal *β*-strand and *α*-helix of the protein are unfolded which interacts with other protein conformations in the solution and initiates dimerization through domain-swapping of the N-terminal *β*-strands. Mutations, which suppress the population of these intermediates should prevent the aggregation of prions. The lifetime of these intermediates is drastically reduced in the disease-resistant huPrP (G127V) revealing a possible role of these conformations in prion aggregation. Experiments can further test the predictions by inducing mutations, which suppress the population of these intermediates to identify key residues that play an important role in the aggregation of prion proteins. Using the native-centric coarse-grained SOP-SC model, we can study only the early stages of domain swapping aggregation in prion protein. The secondary structural transitions that can happen in the later stages of aggregation are not accessible using these models.

## Acknowledgenent

A part of this work is funded by the grants to G.R. from Science and Engineering Research Board (EMR/2016/0013,56) and Nano mission, Department of Science and Technology, India. B.M. acknowledges research fellowship from Indian Institute of Science-Bangalore. The computations are performed using the TUE and Cray XC40 clusters at IISc.

## Graphical TOC Entry

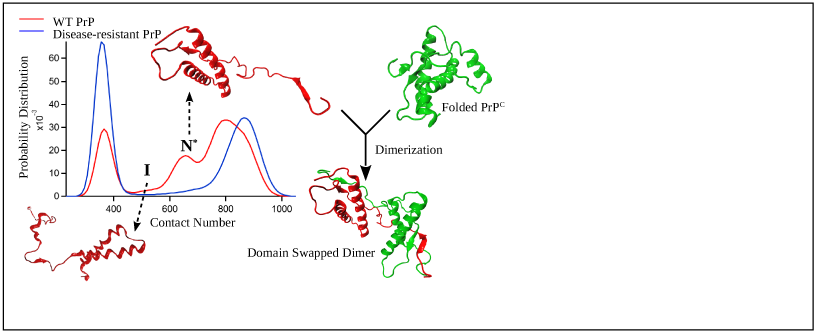

