## Supplementary material for "A Transient Intermediate Populated in Prion Folding Leads to Domain Swapping": SI

### Supporting Information for "A Transient Intermediate Populated in Prion Folding Leads to Domain Swapping"

#### Running header

Supporting Information for "A Transient Intermediate Populated in Prion Folding Leads to Domain Swapping"

### Methods

#### Self Organized Polymer-Side Chain (SOP-SC) Model for Proteins

We have used native-centric SOP-SC model(*1*, *2*) to study the folding thermodynamics of moPrP, wild-type and disease-resistant hPrPs. In SOP-SC model each amino acid is represented by two beads. The backbone atoms of an amino acid are represented by a bead positioned at the center of $C_\alpha$ atom, and the side chain atoms are represented by another bead positioned at the center of mass of the side-chain. The SOP-SC model for moPrP, wild-type hPrP and disease-resistant hPrP are constructed using the structures in the protein data bank (PDB) with PDB ID: 1AG2(*3*), 5YJ5(*4*) and 5YJ4(*4*), respectively. Missing hydrogen atoms are added to the structures using the program visual molecular dynamics (VMD) (*5*) before calculating the centre of mass of the side-chain. The Hamiltonian corresponding to the SOP-SC model is described in terms of bonded ($E_B$), non-bonded ($E_{NB}$) and electrostatics interactions ($E_{el}$). Covalently connected beads interact via a bonded potential ($E_B$). Non-bonded interactions ($E_{NB}$) consist of native (N) and non-native (NN) interactions. Interactions between two beads are considered native, if they are separated by at least three bonds and are within a cut-off distance ($R_c$) in the SOP-SC model of the PDB structure. Any other non-covalent interactions are considered as non-native interactions ($E_{NN}$). Native interactions between neighboring side-chain beads are ignored because of their close proximity in both folded and unfolded states. Electrostatic interactions are present between charged residues and are ignored if charges are present on adjacent side-chain beads. The force-field associated with the SOP-SC model for a protein conformation described by the set of coordinates $\{\mathbf{r}\}$ is given by,

$$E_{CG}(\{\mathbf{r}\}) = E_B + E_{NB}^{N} + E_{NB}^{NN} + E_{el} \tag{S1}$$

The bonds between the beads in the SOP-SC model are modeled using the finite extensible nonlinear elastic (FENE) potential and is given by,

$$E_B = -\sum_{i=1}^{N_B} \frac{k}{2} R_0^2 \log\Big(1 - \frac{(r_i - r_{cry,i})^2}{R_0^2}\Big), \tag{S2}$$

$N_B$ being the total number of bonds present between the covalently linked beads, $r_i$ is the distance between $i^{th}$ pair of beads and $r_{cry,i}$ is the distance between the same $i^{th}$ pair of beads in the SOP-SC PDB structure. The native interactions $E_{NB}^N$, are modeled using a Lennard-Jones type of potential given as,

$$\begin{aligned} E_{NB}^N = &\sum_{i=1}^{N_N^{bb}} \epsilon_h^{bb}\big[\big(\frac{r_{cry,i}}{r_i}\big)^{12} - 2\big(\frac{r_{cry,i}}{r_i}\big)^6\big] + \sum_{i=1}^{N_N^{bs}} \epsilon_h^{bs}\big[\big(\frac{r_{cry,i}}{r_i}\big)^{12} - 2\big(\frac{r_{cry,i}}{r_i}\big)^6\big] \\ &+ \sum_{i=1}^{N_N^{ss}} 0.5 \times 300 k_B \times (0.7 - \epsilon_i^{ss})\big[\big(\frac{r_{cry,i}}{r_i}\big)^{12} - 2\big(\frac{r_{cry,i}}{r_i}\big)^6\big], \end{aligned} \tag{S3}$$

where $N_N^{bb}$, $N_N^{bs}$ and $N_N^{ss}$ represent the total number of native contact pairs present between backbone-backbone, backbone-side chain, and side chain-side chain beads, respectively. $k_B$ is the Boltzmann constant, $r_i$ is the distance between $i^{th}$ pair of beads and $r_{cry,i}$ is the distance between the same $i^{th}$ pair of beads in the SOP-SC PDB structure. $\epsilon_h^{bb}$ and $\epsilon_h^{bs}$ denote the strength of interaction between backbone beads and backbone-side chain beads respectively. Strength of interactions between side-chain beads $\epsilon_i^{ss}$ are obtained from the Betancourt-Thirumalai statistical potential(*6*).

Purely repulsive non-native interactions ($E_{NB}^{NN}$) are modeled as

$$E_{NB}^{NN} = \sum_{i=1}^{N_{NN}} \epsilon_l \big(\frac{\sigma_i}{r_i}\big)^6 + \sum_{i=1}^{N_{ang}^{bb}} \epsilon_l \big(\frac{\sigma^{bb}}{r_i}\big)^6 + \sum_{i=1}^{N_{ang}^{bs}} \epsilon_l \big(\frac{\sigma_i^{bs}}{r_i}\big)^6, \tag{S4}$$

where $N_{NN}$ is the total number of non-native interaction pairs present in the SOP-SC model, $\sigma_i$ is sum of the radii of the beads in $i^{th}$ pair of non-native interactions, $\sigma^{bb}$ is the diameter of backbone bead, and $\sigma^{bs} = \frac{\sigma^{bb}+\sigma_i^{ss}}{2}$ where $\sigma_i^{ss}$ is the diameter of the side chain bead in $i^{th}$

angular interaction between a backbone and side chain beads. The second and third terms in eq. S4 model the bond angle potential between beads separated by two bonds. $N_{ang}^{bb}$ and $N_{ang}^{bs}$ represent the total number of bond angles between the backbone beads and between the backbone-side chain beads. Electrostatic interactions are implemented using a screened coulomb potential given as

$$E_{el} = \sum_{i=1}^{N_c-1} \sum_{j=i+1}^{N_c} \frac{q_i q_j exp(-\kappa r_{ij})}{\epsilon r_{ij}}, \tag{S5}$$

where $N_c$ is the number of charged residues in the protein, $r_{ij}$ is the distance between the charged side chains $i$ and $j$, $q_i$ and $q_j$ are the point charges measured in units of electron charge placed on the centers of the charged side chain beads $i$ and $j$, respectively. At neutral pH, $q_i$ is considered +1 for positively charged residues, and -1 for negatively charged residues. The inverse Debye length, $\kappa$, accounts for the presence of a monovalent salt of 10mM concentration. For implicit solvent description, usual range of dielectric constants are from 2 to 20(*7*). In the simulations, we considered the dielectric constant of the medium as $\epsilon$ = 10 $\epsilon_0$, where $\epsilon_0$ is the vacuum permittivity. Values of the parameters used to describe the SOP-SC energy functions and side-chain radii are listed in Table S1, Table S2 and Table S3, respectively.

#### Symmetric Go potential to mimic inter-protein interactions

To mimic the interactions between two prion protein chains, we employed symmetrized Go-type hamiltonians(*8*, *9*), where native contacts present in a monomer chain is used to define the additional inter-protein interactions present in the dimeric system. Fig. S9 depicts the essential features of symmetric Go-type potential where for each native interaction present between residues $i$ and $j$ of chain A, there are three additional native intercations present between $i'$ of chain B and $j$ of chain A, $i$ of chain A and $j'$ of chain B and finally between $i'$ and $j'$ of chain B. The operative force field in the dimeric prion systems (monomer $M_1$ and

$M_2$) can be described as,

$$E_{M_1,M_2}(\{\mathbf{r}\}) = E_B(M_1) + E_B(M_2) + E_{NB}^N(M_1) + E_{NB}^N(M_2) \\ + E_{NB}^{NN}(M_1) + E_{NB}^{NN}(M_2) + E_{NB}^N(M_1, M_2) + E_{NB}^{NN}(M_1, M_2), \quad \text{(S6)}$$

where $E_B(M_1)$ and $E_B(M_2)$ represent bonded interactions in the prion monomers $M_1$ and $M_2$, $E_{NB}^N(M_1)$ and $E_{NB}^N(M_2)$ represent non-bonded native interactions in $M_1$ and $M_2$, and $E_{NB}^{NN}(M_1)$ and $E_{NB}^{NN}(M_2)$ represent non-bonded non-native interactions in $M_1$ and $M_2$. The terms $E_{NB}^N(M_1, M_2)$ and $E_{NB}^{NN}(M_1, M_2)$ in eq. S6 represent the inter native contacts and non-native contacts present between $M_1$ and $M_2$, respectively.

The inter native interactions between the monomers $M_1$ and $M_2$ is given by

$$E_{NB}^N(M_1, M_2) = \sum_{i=1}^{N_N^{bb}(M_1,M_2)} \epsilon_h^{bb}\big[\big(\frac{r_{cry,i}}{r_i}\big)^{12} - 2\big(\frac{r_{cry,i}}{r_i}\big)^6\big] + \sum_{i=1}^{N_N^{bs}(M_1,M_2)} \epsilon_h^{bs}\big[\big(\frac{r_{cry,i}}{r_i}\big)^{12} - 2\big(\frac{r_{cry,i}}{r_i}\big)^6\big] \\ + \sum_{i=1}^{N_N^{ss}(M_1,M_2)} 0.5 \times 300k_B \times (0.7 - \epsilon_i^{ss})\big[\big(\frac{r_{cry,i}}{r_i}\big)^{12} - 2\big(\frac{r_{cry,i}}{r_i}\big)^6\big], \quad \text{(S7)}$$

where $N_N^{bb}(M_1, M_2)$, $N_N^{bs}(M_1, M_2)$ and $N_N^{ss}(M_1, M_2)$ represent inter native backbone-backbone, backbone-side chain, and side chain-side chain interactions, respectively. The inter non-native interactions between $M_1$ and $M_2$ are given by

$$E_{NB}^{NN}(M_1, M_2) = \sum_{i=1}^{N_{NN}(M_1,M_2)} \epsilon_l\big(\frac{\sigma_i}{r_i}\big)^6, \quad \text{(S8)}$$

where $N_{NN}(M_1, M_2)$ is the total number of inter non-native interactions present between the beads of $M_1$ and $M_2$. The necessary parameters are listed in Table S1 and Table S2.

#### Simulations

We employed low friction Langevin dynamics simulations to effectively sample protein's conformational space and calculated important thermodynamic properties. The equation of motion for a bead with position coordinates $\vec{r_i}$ is depicted as,

$$m\ddot{\vec{r_i}} = -\zeta\dot{\vec{r_i}} + \vec{F_c} + \vec{\Gamma}, \tag{S9}$$

, where m is the mass of the respective protein bead, $\zeta$ is the friction co-efficient, $\vec{F_c} = -\frac{\partial E_{CG}(\{\mathbf{r}\},0,t)}{\partial \vec{r_i}}$, $\vec{\Gamma}$ is the random force with a white noise spectrum. The autocorrelation function of the random force in the discretised form is given by $\langle\Gamma(t)\ \Gamma(t+nh)\rangle = \frac{2\zeta k_B T}{h}\delta_{0,n}$, where $n = 0, 1, ...$ and $\delta_{0,n}$ is the Kronecker delta function. Velocity Verlet algorithm(*10*, *11*) is used to integrate the equation of motion. We used $\zeta = 0.05\ m/\tau_L$ and time step, $h = 0.005\ \tau_L$, where $\tau_L$ is the unit of time used to advance the simulation.

To probe the folding kinetics of proteins, we used brownian dynamics simulations where equations of motions are integrated using the Ermak-McCammon algorithm(*12*),

$$\vec{r_i(t+h)} = \vec{r_i(t)} + \frac{h}{\zeta}\vec{F_c} + \vec{\Gamma}. \tag{S10}$$

$\vec{\Gamma}$ is the random force with mean $\langle\Gamma(h)\rangle = 0$, and variance $\langle\Gamma(h)^2\rangle = \frac{2k_BTh}{\zeta}$. The friction coefficient $\zeta = 31.9\ m/\tau_H$ is chosen close to that of water and integrating time step $h$ is set to $0.05\tau_H$. In the simulations, we chose the unit of length $a = 1$ Å, energy $\epsilon = 1$ kcal/mol, and mass $m = 1.8 * 10^{-22}$ g. Unit of time in Langevin dynamics simulations is $\tau_L = \sqrt{ma^2/\epsilon} = 0.51$ ps, while time step $\tau_H$ in Brownian dynamics simulations is $\tau_H \approx \frac{\zeta_H a^2}{k_BT} = \frac{(\zeta_H\tau_L/m)\epsilon}{k_BT}\tau_L \approx 53$ ps. We used weighted histogram analysis method (WHAM)(*13*) to compute average thermodynamic properties from Langevin dynamics simulations performed at a range of temperatures. The detailed description of the WHAM equations to compute equilibrium properties is reported elsewhere(*14*).

#### Atomistic Simulations

We transformed the coarse-grained domain swapped structure of moPrP to an all atom description using CG Builder module (*15*, *16*) in VMD. The dimeric system is solvated in a cubic box of length of 90 Å containing ≈ 22000 TIP3P(*17*) water molecules. The simulation box is neutralized with NaCl. All atom simulations are carried out using NAMD molecular dynamics package(*18*). CHARMM22 force-field parameters (*19*, *20*) are used for protein. Simulations are run in the $NPT$ ensemble with pressure $P$ = 1 atm and temperature $T$ = 300 K using Nosé-Hoover Langevin piston with a barostat oscillation time 200 fs(*21*, *22*). Periodic boundary conditions are applied along $x$, $y$ and $z$- directions. Particle Mesh Ewald (PME) method(*23*) is used to calculate long range electrostatics interactions with a grid size of 1 Å. The distance cutoff for non-bonded interactions for both Coulomb and van der Waals potentials is 12 Å. The simulation is advanced using a time step of 2 fs. All the bonds involving hydrogen atoms are constrained using RATTLE algorithm(*24*). Conjugate gradient algorithm is used to minimize the initial structures. After minimization, we performed MD simulations on three independent prion dimeric systems for at least 500 ns each.

Table S1: Energy function parameters for the SOP-SC model

| Parameters | Values |
|---|---|
| $R_o$ | 2.0 |
| $k$ | 20 kcal/(mol. Å$^2$) |
| $R_c$ | 8 Å |
| $r_0$ | 2 Å |
| $\epsilon_h^{bb}$ | 0.45 kcal/mol |
| $\epsilon_h^{bs}$ | 0.45 kcal/mol |
| $\epsilon_l$ | 1.0 kcal/mol |
| $\sigma^{bb}$ | 3.8 Å |

Table S2: SOP-SC model parameters for prion protein variants

| Parameters | 1AG2 | 5YJ4 | 5YJ5 |
|---|---|---|---|
| $N_B$ | 205 | 213 | 213 |
| $N_N^{bb}$ | 250 | 246 | 242 |
| $N_N^{bs}$ | 709 | 710 | 682 |
| $N_N^{ss}$ | 300 | 291 | 283 |
| $N_N^{bb}(M_1, M_2)$ | 500 | - | - |
| $N_N^{bs}(M_1, M_2)$ | 1418 | - | - |
| $N_N^{ss}(M_1, M_2)$ | 600 | - | - |
| $N_{NN}$ | 19244 | 20908 | 20948 |
| $N_{NN}(M_1, M_2)$ | 39918 | - | - |
| $N_{ang}^{bb}$ | 101 | 105 | 105 |
| $N_{ang}^{bs}$ | 204 | 212 | 212 |
| $N_c$ | 22 | 25 | 25 |

Table S3: Side-chain radii of amino acids

| Residue | Radius (Å) |
|---|---|
| Gly | 0.5 |
| Ala | 2.52 |
| Val | 2.93 |
| Leu | 3.09 |
| Ile | 3.09 |
| Met | 3.09 |
| Phe | 3.18 |
| Pro | 2.78 |
| Ser | 2.59 |
| Thr | 2.81 |
| Asn | 2.84 |
| Gln | 3.01 |
| Tyr | 3.23 |
| Trp | 3.39 |
| Asp | 2.79 |
| Glu | 2.96 |
| Hsd | 3.04 |
| Lys | 3.18 |
| Arg | 3.28 |
| Cys | 2.74 |

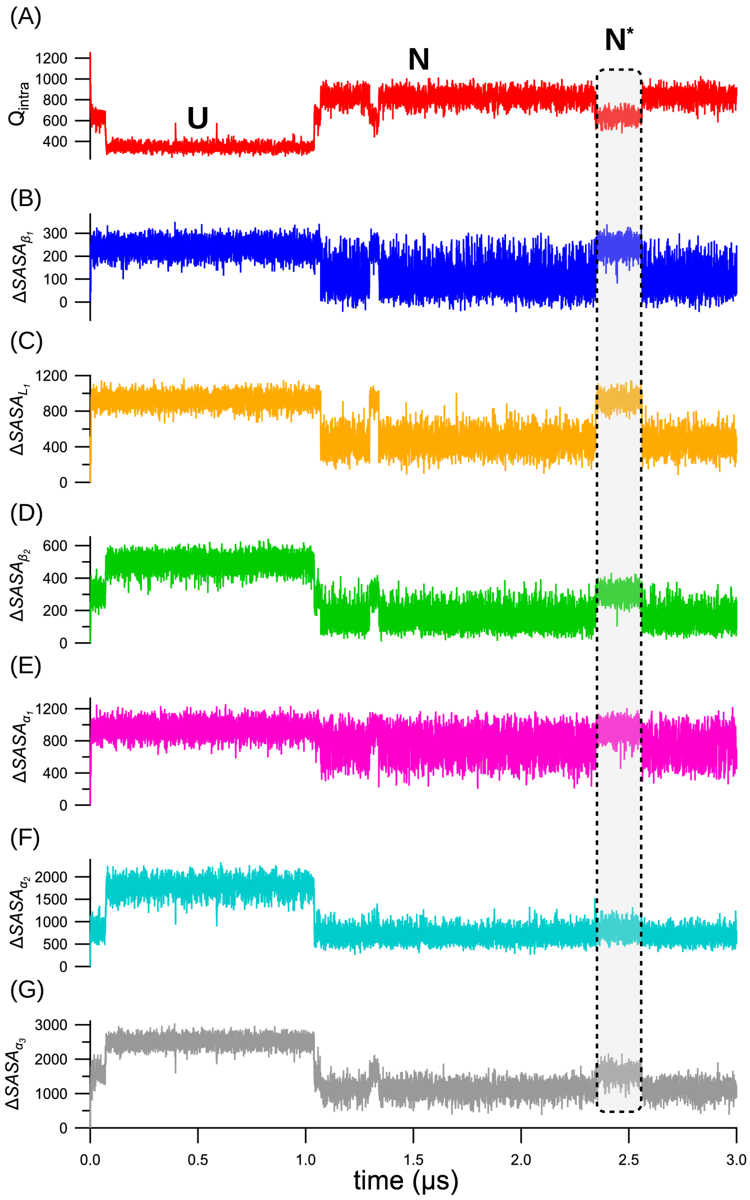


Figure S1: Time evolution of native contacts $Q_{intra}$ and change in the solvent accessible surface area (SASA) of different secondary structural elements (SSEs) relative to the native folded state, $\Delta SASA_{SSE}$ (in Å$^2$ unit) in moPrP at $T_M$. (A) At $T_M$, the protein samples the folded state $N$ ($Q_{intra} \approx 900$), unfolded state $U$ ($Q_{intra} \approx 350$) and intermediate state $N^*$ ($Q_{intra} \approx 600$). $N^*$ state is highlighted using a rectangular shaded area. (B), (C), (D), (E), (F), and (G) shows $\Delta SASA_{SSE}$ for SSEs $\beta_1$, $L_1$, $\beta_2$, $\alpha_1$, $\alpha_1$ and $\alpha_3$, respectively for the trajectory shown in (A). In the $N^*$ state, $\beta_1$, $\alpha_1$ and $L_1$ undergo complete unfolding. Structural perturbation is also observed in the folded core $\beta_2 - \alpha_2 - \alpha_3$ of $N^*$ state. The SSE $\alpha_2$ retains its folded form while $\beta_2$ and $\alpha_3$ undergo partial unfolding resulting in an increase in their respective $\Delta SASA_{SSE}$s.

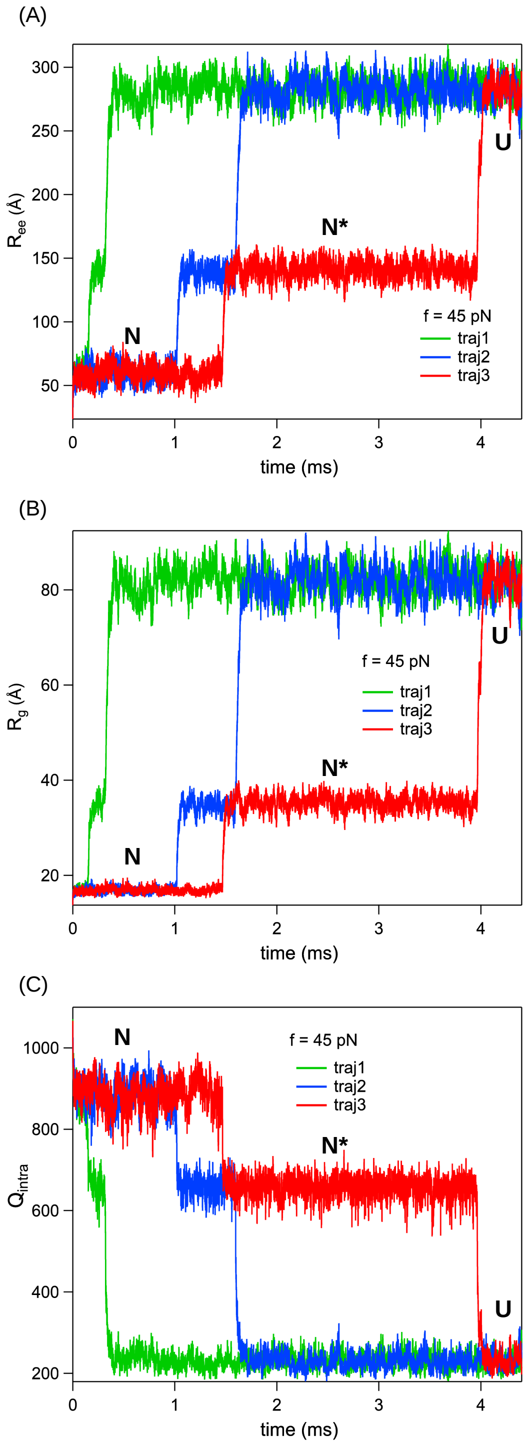


Figure S2: Time evolution of (A) $R_{ee}$ (B) $R_g$ and (C) $Q_{intra}$ during force induced unfolding of moPrP at $T = 300$ K and $f = 45$ pN for three independent trajectories are shown in red, blue and green. An intermediate state $N^*$ is populated at $Q_{intra} \approx 700$ and $R_g \approx 38$ Å in addition to the folded ($N$) and unfolded states ($U$).

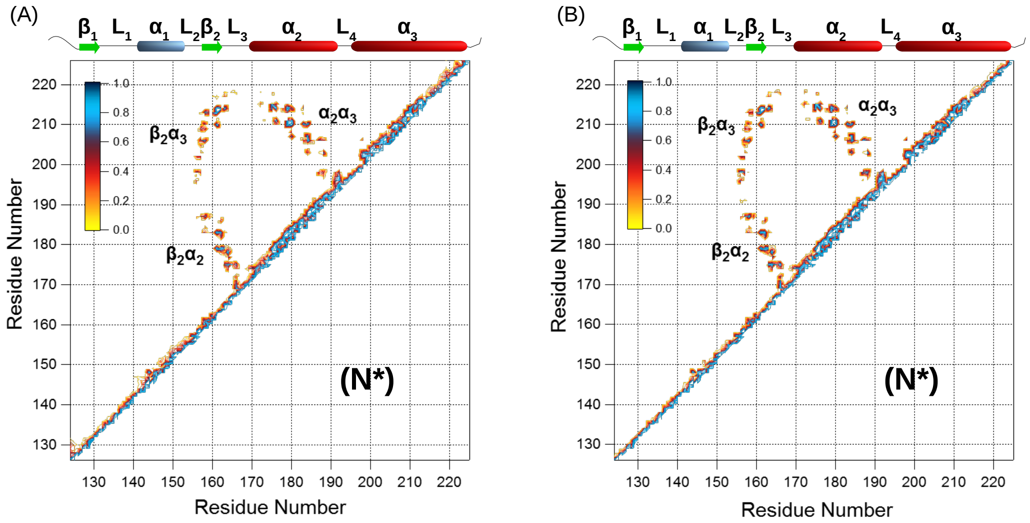


Figure S3: Contact map of the $N^*$ states of moPrP obtained from (A) equilibrium and (B) force induced pulling simulations at $f = 45$ pN. Strong resemblance between the two contact maps reveals that the tertiary contacts present in both the $N^*$ states are similar.

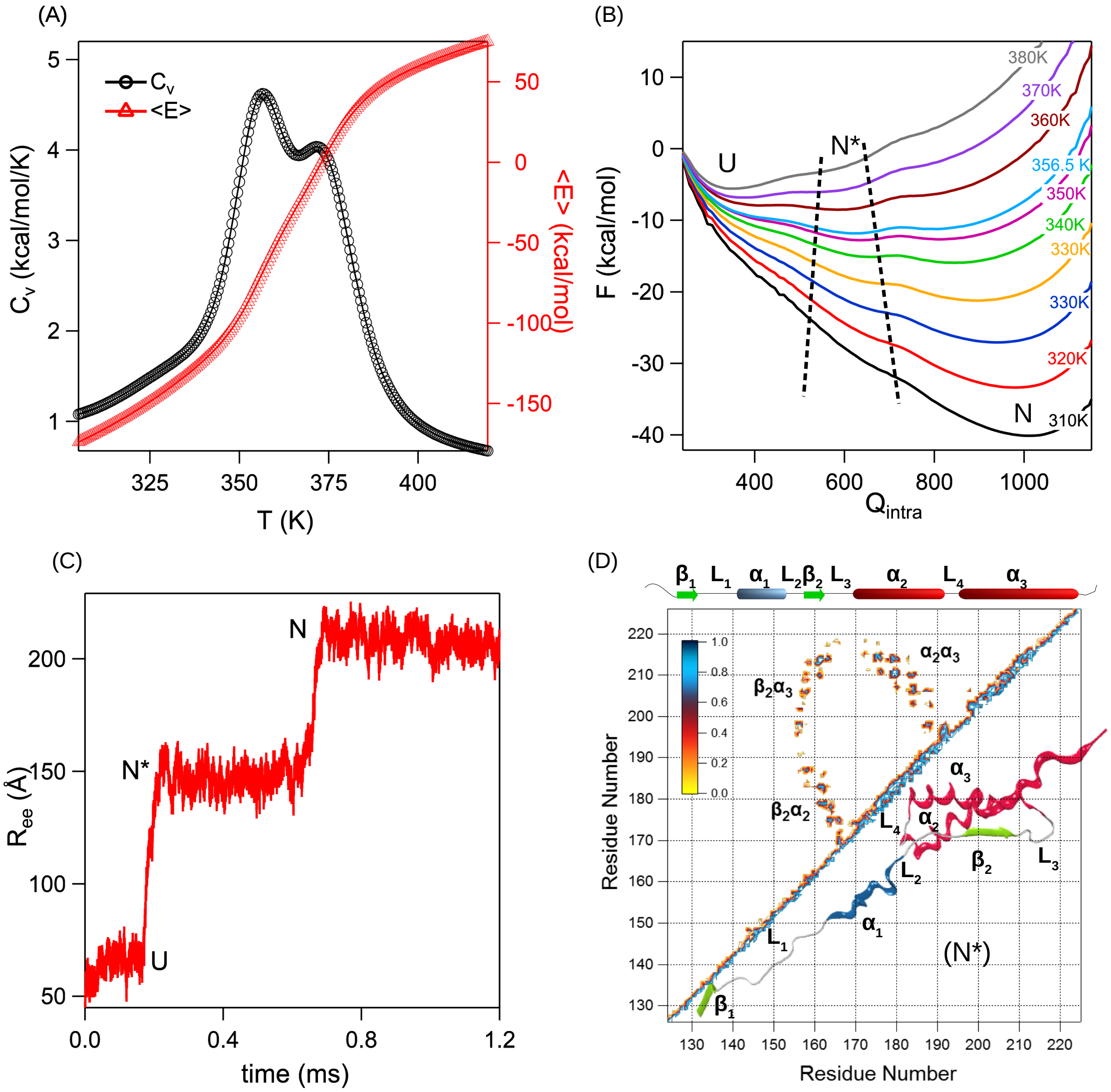


Figure S4: (A) Specific heat capacity $C_v$ and average potential energy $\langle E \rangle$ are plotted as a function of temperature $T$ for moPrP with a disulfide bond (Cys179-Cys214). $C_v$ indicates a three state folding-unfolding process with a well populated intermediate state ($N^*$). (B) Free energy of folding $\Delta F$ projected onto $Q_{intra}$ for different $T$s ranging from 310 K to 380 K shows the population of $N^*$ state. (C) Mechanical unfolding of moPrP at a constant force $f = 60$ pN in presence of a disulfide bond. $R_{ee}$ is plotted as a function of simulation time. Three state unfolding is observed with a well populated intermediate $N^*$. (D) Contact map of $N^*$ reveals a structured core comprised of $\beta_2 - \alpha_2 - \alpha_3$ and a disordered segment $\beta_1 - \alpha_1$.

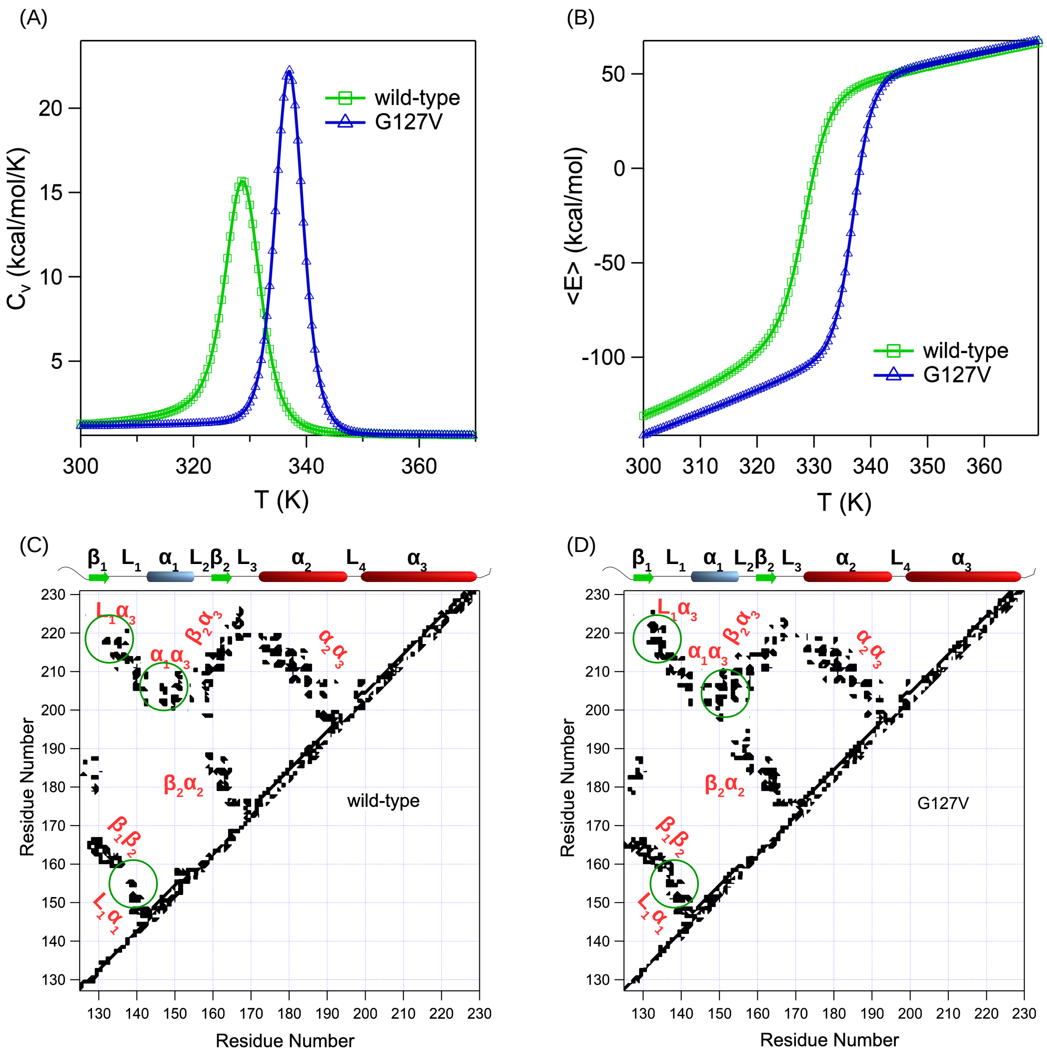


Figure S5: Effect of single point mutation (G127V) on thermodynamic stability of hPrPs (A) Specific heat capacity $C_v$ and (B) average potential energy $\langle E \rangle$ plotted as a function of temperature $T$ indicate two state folding of hPrP. Disease-resistant hPrP is thermally more stable than wild-type hPrP. Contact map of (C) wild-type (PDB ID: 5YJ4) and (D) disease-resistant hPrP (PDB ID: 5YJ5) reveal an increased number of contacts between $\alpha_1$ and $\alpha_3$, $L_1$ and $\alpha_3$, and, $L_1$ and $\alpha_1$ in the disease-resistant hPrP, resulting in a relatively more compact folded state.

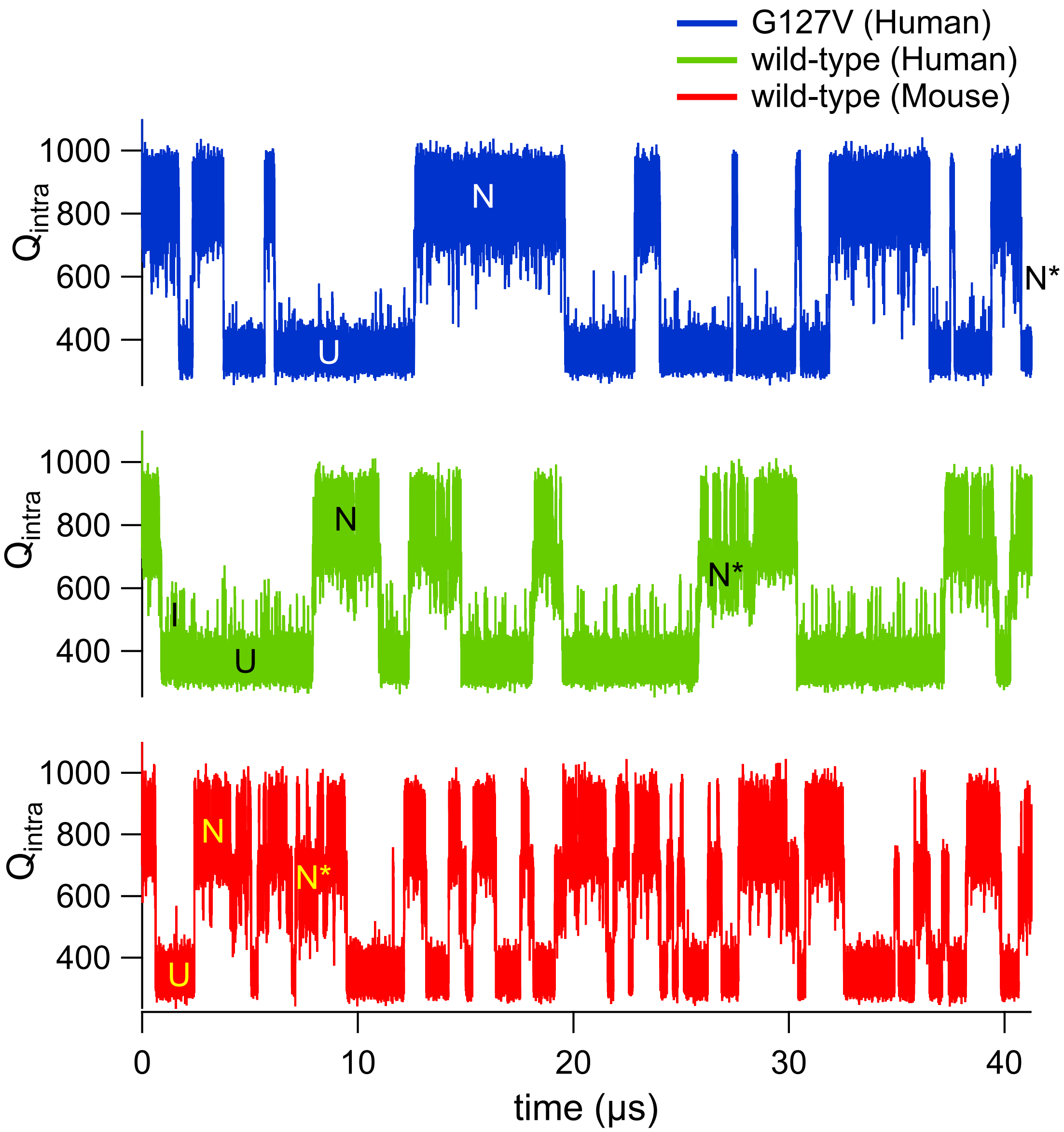


Figure S6: Time evolution of $Q_{intra}$ at respective $T_M$s for wild-type moPrP (in red), wild-type hPrP (in green) and disease-resistant hPrP (in blue). Metastable intermediate states ($N^*$ and $I$) are observed in the trajectories of wild type prion proteins, while for disease-resistant hPrP, fleeting transitions to the intermediate states are observed.

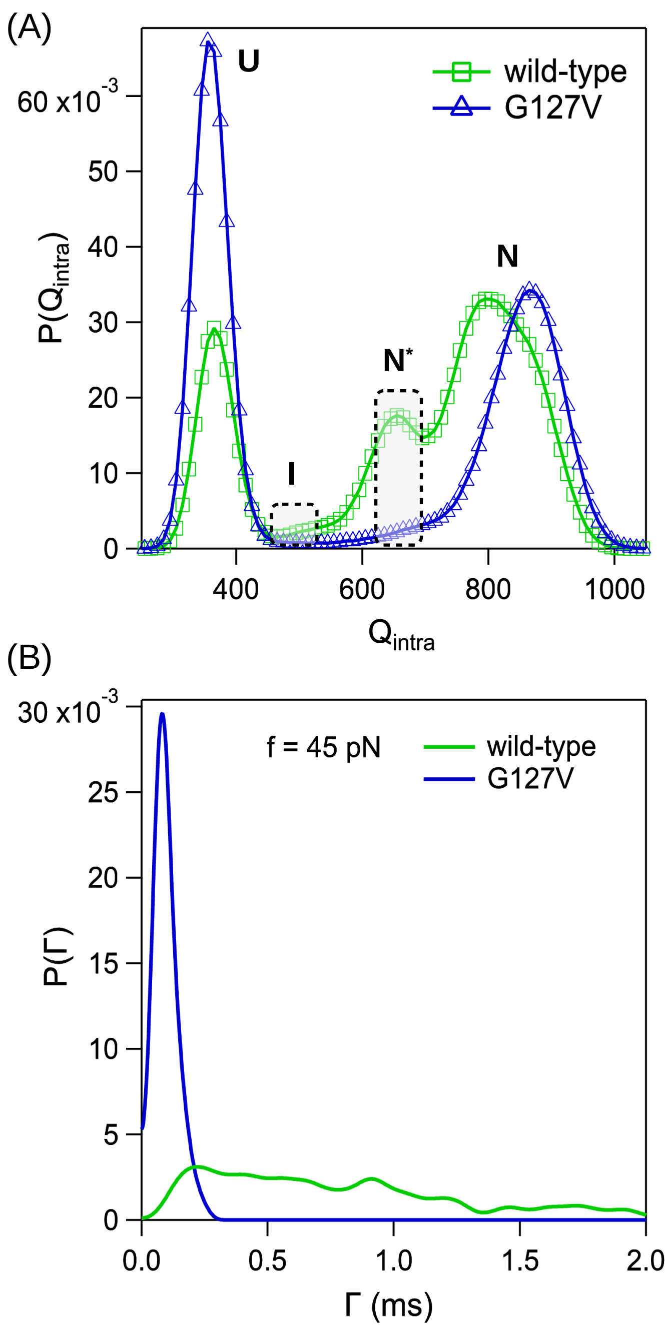


Figure S7: (A) Probability distribution of $Q_{intra}$ ($P(Q_{intra})$) is plotted for wild-type and disease-resistant hPrPs at their respective $T_M$s. In addition to $N$ ($Q_{intra} \approx 900$) and $U$ ($Q_{intra} \approx 400$) states, two intermediate states, $I$ ($Q_{intra} \approx 500$) and $N^*$ ($Q_{intra} \approx 650$), are populated in wild-type prion. State $I$ is absent in disease-resistant prion, and stability of $N^*$ is drastically reduced. (B) Probability distribution of $N^*$ state lifetime $\Gamma$ ($P(\Gamma)$) computed from 350 constant force pulling simulations performed at $T = 300$ K and $f = 45$ pN show that $P(\Gamma)$ is narrow with an average lifetime of 0.1 ms in disease-resistant prion while $P(\Gamma)$ of $N^*$ state in the wild-type protein is broad with an average lifetime of 0.7 ms.

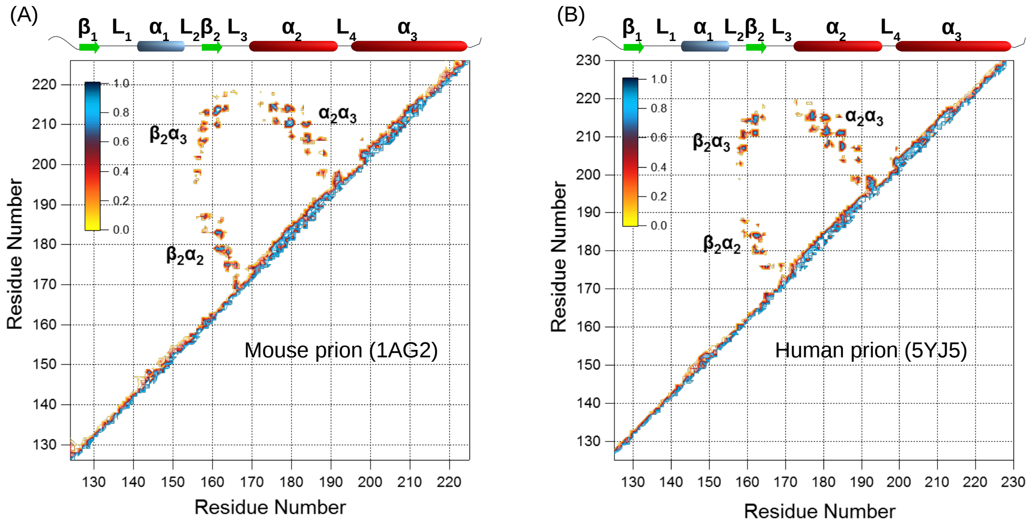


Figure S8: Contact maps of $N^*$ state in (A) moPrP (PDB ID: 1AG2) and (B) hPrP (PDB ID: 5YJ5). Tertiary contacts are present between the SSEs $\beta_2$, $\alpha_2$ and $\alpha_3$, while contacts are absent in $\beta_1 - L_1 - \alpha_1$ segment. This suggests a possibility of a common $N^*$ structure across different mammalian species.

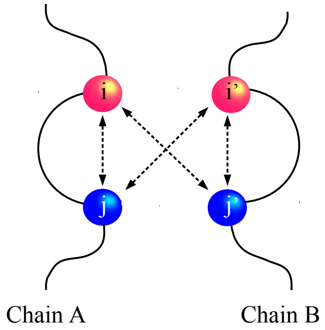


Figure S9: Symmetric Go model potential between two peptide chains A and B. Any native interaction present between residues $i$ and $j$ in chain A, will also be present between $i$ of chain A and $j'$ of chain B; $j$ of chain A and $i'$ of chain B; and between $i'$ and $j'$ of chain B.

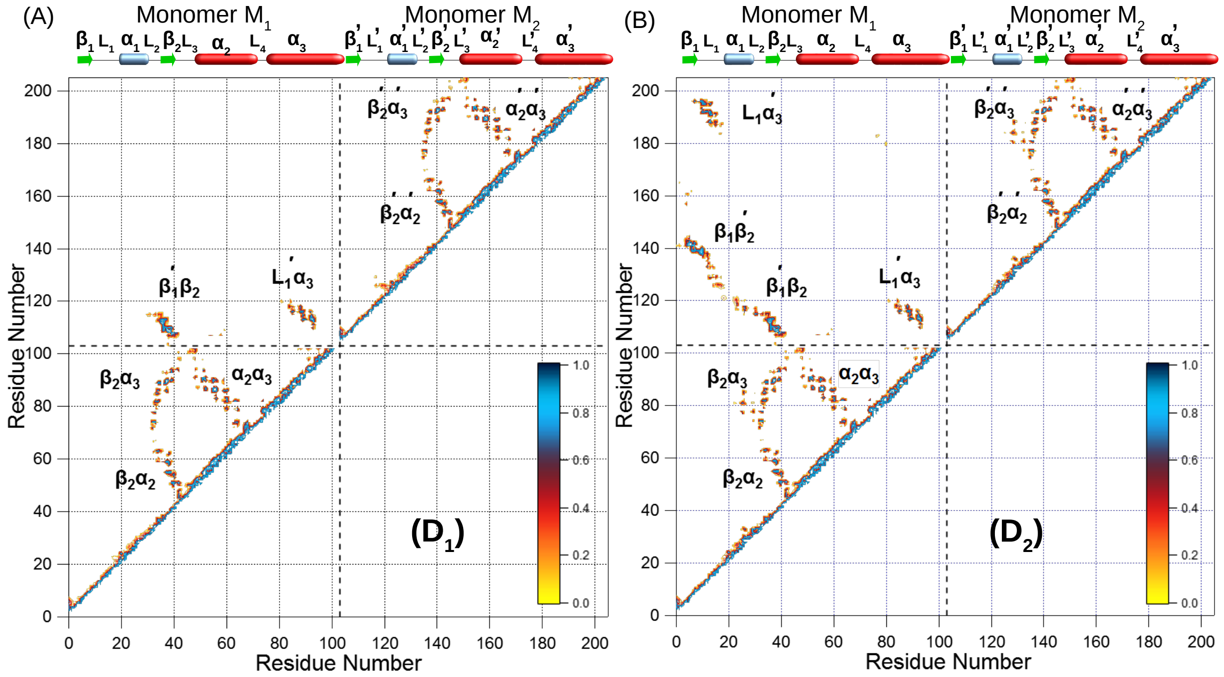


Figure S10: Contact map of (A) $D_1$ and (B) $D_2$ states show a partial and fully formed domain swapped structures. In the $D_1$ state, $\beta_1'$ in $M_2$ domain swaps and interacts with $\beta_2$ in $M_1$, whereas $\beta_1$ in $M_1$ has not domain-swapped to interact with $\beta_2'$ in $M_2$. In $D_2$ complete domain swapping is observed.

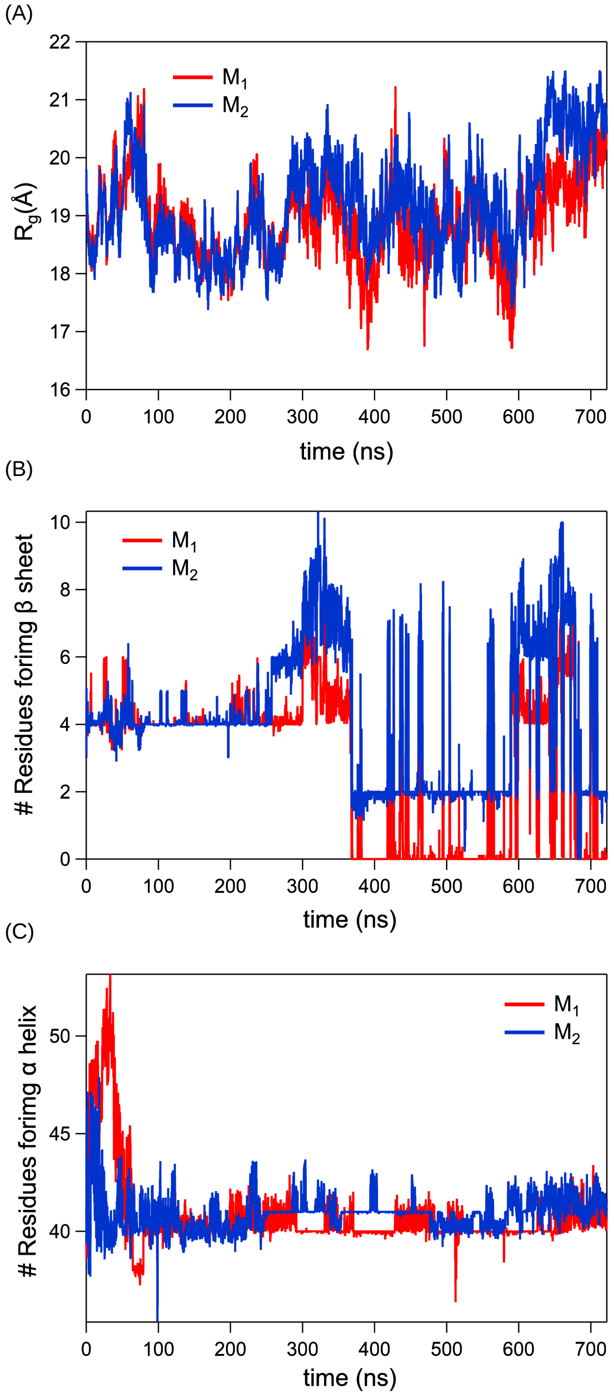


Figure S11: Time evolution of (A) $R_g$, (B) number of residues forming $\beta$-sheet ($N_\beta$) and (C) number of residues forming $\alpha$-helix ($N_\alpha$) in monomer $M_1$ (in red) and monomer $M_2$ (in blue) in atomistic NPT simulations of moPrP dimer performed at $T = 300$ K. The dimeric domain swapped structure of moPrP obtained from coarse-grained simulations is stable over a time period of 700 ns.
